# Rapid Turnover of Life-Cycle-Related Genes in the Brown Algae

**DOI:** 10.1101/290809

**Authors:** A.P. Lipinska, M.L. Serrano-Serrano, Akira F. Peters, K. Kogame, J Mark Cock, Susana M. Coelho

## Abstract

**Background:** Sexual life cycles in eukaryotes involve a cyclic alternation between haploid and diploid phases. While most animals possess a diploid life cycle, plants and algae alternate between multicellular haploid (gametophyte) and diploid (sporophyte) generations. In many algae, gametophytes and sporophytes are independent and free living, and may present dramatic phenotypic differences. The same shared genome can therefore be subject to different, even conflicting, selection pressures in each of the life cycle generations. Here, we have analysed the nature and extent of genome-wide generation-biased gene expression in four species of brown algae with contrasting levels of dimorphism between life cycle generations, in order to assess the potential role of generation-specific selection in shaping patterns of gene expression and divergence.

**Results:** We show that the proportion of the transcriptome that is generation-biased is associated with the level of phenotypic dimorphism between the life cycle stages. Importantly, our data reveals a remarkably high turnover rate for life-cycle-related gene sets across the brown algae and highlights the importance not only of co-option of regulatory programs from one generation to the other but also a key role for newly emerged, lineage-specific genes in the evolution of the gametophyte and sporophyte developmental programs in this major eukaryotic group. Moreover, we show that generation-biased genes display distinct evolutionary modes, with gametophyte-biased genes evolving rapidly at the coding sequence level whereas sporophyte-biased genes exhibit changes in their patterns of expression.

**Conclusion:** Our analysis uncovers the characteristics, expression patterns and evolution of generation-biased genes and underline the selective forces that shape this previously underappreciated source of phenotypic diversity.

## BACKGROUND

As a consequence of sexual reproduction, the vast majority of eukaryotes have life cycles involving an alternation between haploid and diploid phases [1,2]. The proportion of the life cycle spent in each phase varies dramatically depending on the species. In organisms with haplontic cycles, mitosis only occurs in the haploid stage. Haploid mitosis may lead to asexual (clonal) reproduction such as the case of *Chlamydomonas*, or involve somatic growth and cellular differentiation such as in *Volvox*. In these organisms, the zygote undergoes meiosis immediately after syngamy without undergoing any mitotic divisions. Conversely, in diplontic life cycles, mitosis only occurs during the diploid phase, and meiosis takes place immediately before gamete formation. Diploid mitosis leads to asexual reproduction in unicellular lineages (e.g. diatoms) and to somatic growth and differentiation in multicellular organisms such as Metazoans. Finally, in organisms with haploid-diploid life cycles, mitotic cell divisions occur during both the haploid and diploid phases. In land plants and some algae, these mitotic divisions can lead to the development of two distinct multicellular organisms, one haploid and the other diploid. The haploid organism is generally referred to as the gametophyte, because it produces gametes, and the diploid organism as the sporophyte, because it produces spores. Note, however, that the gametophyte and sporophyte developmental programs are not absolutely linked to ploidy because ploidy and life cycle generation have been shown to be uncoupled during variant life cycles [3,4]. The gametophyte and sporophyte should therefore be thought of as genetically-controlled developmental programs that are coordinated with, but not absolutely linked to, life cycle progression.

The evolutionary advantages of life cycles with dominant haploid, dominant diploid, or alternation between two phases have been subject to extensive theoretical work [5–10]. Models exploring the evolution of haploidy and diploidy assume an alternation of generations with free-living haploid and diploid phases, where expanding one phase reduces the other phase. These models predict that purging of deleterious mutations favours expansion of the haploid phase when recombination is rare, but that diploids are favoured when recombination is common because they mask mutations from selection [6,11]. In contrast, niche differentiation between haploids and diploids may favour the maintenance of biphasic life cycles, in which development occurs in both phases [12]. For instance, gametophytes have been shown to exploit low-resource environments more efficiently whereas sporophytes are more vigorous when resources are abundant [13]. The interplay between genetic and ecological factors has been recently explored [7], in a model that assumes different effects of mutations in haploids and diploids and competition between individuals within a generation. The model predicts that temporal variations in ecological niches stabilize alternation of generations. Empirical support for these models has come from the brown alga *Ectocarpus* sp., where dimorphism between generations has been linked to the occupation of different spatio-temporal niches [14].

In organisms with complex life cycles, an allele may be relatively beneficial when expressed in one generation but deleterious when expressed in the other generation (generation-antagonism), and in this case selection acts in opposite directions in haploids and diploids [7,15]. With this type of generation-dependent antagonistic selection, evolution favours the expansion of whichever generation gains the greatest fitness advantage, on average, from the conflicting selection pressures [15]. Generation antagonism is expected to be particularly relevant in multicellular species where there is alternation of generations with morphologically dissimilar gametophytes and sporophytes, as in the case of many plants and algae. When fitness optima differ between the gametophyte and sporophyte generations for a shared trait, dimorphism can allow each generation to express its optimum trait phenotype. Accordingly, the evolution of generation-biased gene expression may be one mechanism that could help to resolve this intra-locus ‘generation’ conflict, in a similar manner to mechanisms that resolve sexual antagonism [16,17]. Another potential solution to resolve generation-conflict is gene duplication, followed by divergence of the two loci towards distinct optima corresponding to each of the two generations. An equivalent process has been shown to be important in the generation of sex-biased gene expression [17,18]. While the role of sexual selection in shaping phenotypic diversity and in driving patterns of evolution of gene expression has been studied extensively (e.g. [19,20]), we have remained so far largely ignorant about the relationships between generation-biased selection, generation-biased gene expression and phenotypic differentiation.

The brown algae (Phaeophyceae) are a group of complex multicellular eukaryotes that diverged from plants and animals more than a billion years. Brown algal life cycles are extraordinarily diverse, exhibiting a broad range of variation in terms of the relative complexities of the gametophyte and sporophyte generations [21]. Here, we selected two pairs of brown algal species from the orders Ectocarpales and Laminariales, which diverged about 95 Mya [22], to trace the evolutionary history of generation-biased gene expression in the brown algal lineage. The selected species exhibit markedly different levels of dimorphism between life cycle generations: the Laminariales species *Macrocystis pyrifera* and *Saccharina japonica* have complex sporophyte but highly reduced gametophyte generations, whereas the Ectocarpales species include *Scytosiphon lomentaria,* which has a reduced sporophyte but a morphologically complex gametophyte, and *Ectocarpus* sp. which has gametophyte and sporophyte generations of similar complexity. We show that a large proportion of the transcriptome of brown algae exhibit generation-biased expression, and that the set of life-cycle-biased genes turns over extremely rapidly during evolution due to a combination of two processes: *de novo* birth of genes with generation-biased expression and gain/loss of generation-biased expression by orthologous loci. Our results uncover the characteristics, expression patterns and evolution of generation-biased genes and underline the selective forces that shape this previously underappreciated source of phenotypic diversity.

## RESULTS

### Measures of phenotypic differentiation between gametophyte and sporophyte generations

The number of different cell types in each generation and the ratios of the sizes of the gametophyte and the sporophyte at maturity were used as proxies to assess the degree of morphological complexity and the level of phenotypic dimorphism between life cycle generations in the four brown algal species studied (Table S1). Using these parameters, the Laminariales species *M. pyrifera* and *S. japonica* exhibited the highest level of phenotypic differentiation between generations, with the sporophyte being more complex than the gametophyte, both in terms of number of cell types and in terms of the size. As far as the Ectocarpales species were concerned, dimorphism between generations were also marked in *S. lomentaria*, but with the gametophyte being more complex than the sporophyte. *Ectocarpus* sp. exhibited the lowest level of differentiation between the gametophyte and sporophyte generation (Table S1; Figure S1).

### Patterns of generation-biased gene expression in gametophytes and sporophytes

We used DEseq2 to compare patterns of gene expression in gametophytes and sporophytes for each of the four brown algal species. The highest level of generation-biased gene expression was detected in *M. pyrifera*, where 46% of the transcriptome was generation-biased. Generation-biased genes represented 37%, 36% and 35% of the transcriptomes of *S. lomentaria, S. japonica* and *Ectocarpus* sp., respectively (Figure 1A). In general, more transcripts were gametophyte-biased than sporophyte-biased. This difference was most marked in *S. lomentaria*, where almost twice as many genes were gametophyte-biased (27% of the transcriptome) than were sporophyte-biased. In both *Ectocarpus* sp. and *S. lomentaria*, the fraction of sporophyte-biased transcripts was relatively low (12%) but the proportion was higher in species that have a more conspicuous sporophyte generation (i.e., both Laminariales), with 19-22% of the transcriptome being sporophyte-biased (Figure 1A).

**Figure 1.**
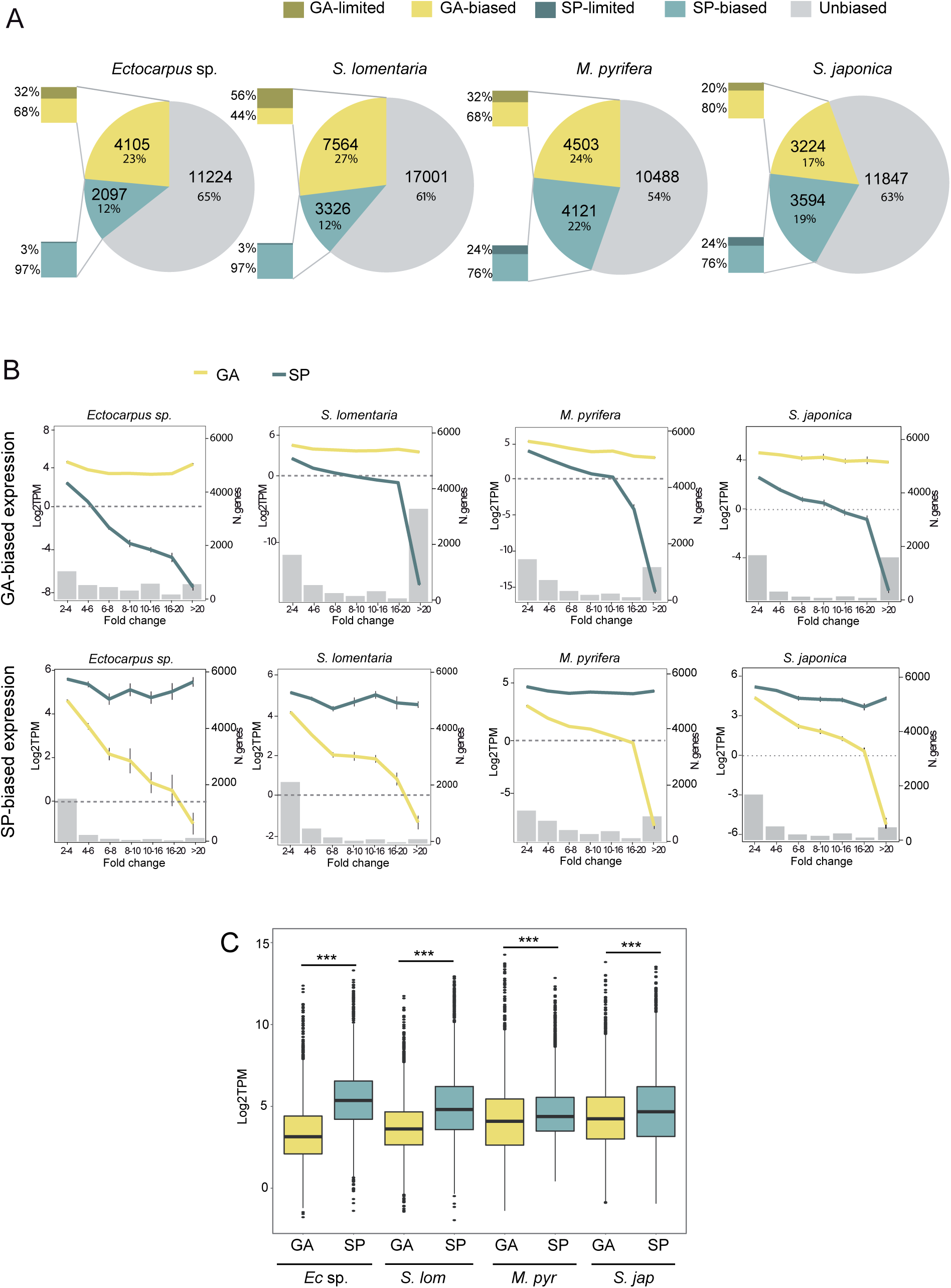
Generation-biased gene expression across the four-brown algal species. **(A)** Proportion of unbiased, gametophyte- and sporophyte-biased genes across the four-studied species. Bar inserts represent the proportion of generation-specific genes among the generation-biased genes in each species. **(B)** Mean gene expression levels (log2TPM) at several degrees of generation-bias (fold change, FC, represented by grey histograms) for gametophyte-biased (yellow) and sporophyte-biased (blue) genes in the four-studied species. The number of genes in each category of FC are represented in the right side of the graph. Error bars represent standard errors. GA: gametophyte; SP: sporophyte. **(C)** Boxplot showing the mean expression levels (log2TPM) of gametophyte- and sporophyte-biased genes.

Generation-biased genes were defined as being generation-limited when the TPM for one of the two generations was below the 5^th^ percentile (see Methods) (Figure 1A). Between 35% and 59% of the generation-biased genes were generation-limited, depending on the species. The proportion of generation-limited genes was larger for the gametophyte than for the sporophyte generation in *Ectocarpus* sp. and *S. lomentaria*. This trend was particularly marked for *S. lomentaria* (which has a dominant gametophyte generation) where more than half of the gametophyte-biased genes were gametophyte-limited. In contrast, in the two Laminariales species, similar proportions of generation-limited genes were observed in both generations (Figure 1A).

To examine the relationship between degree of generation-biased expression and transcript abundance (expression level), the generation-biased genes were grouped according to the fold change (FC) difference between gametophyte and sporophyte samples, and the mean expression levels in gametophytes and sporophytes (log_2_TPM) were plotted for each group (Figure 1B). This analysis indicated that, overall, the most marked levels of generation-biased expression (high fold changes) were the result of down-regulation of genes in the generation where they were expressed more weakly, rather than strong up-regulation in the generation where they were expressed more strongly. However, for gametophyte-biased genes, the expression in sporophytes reached the lower threshold (about log_2_TPM<0) much faster than the expression of sporophyte-biased genes in gametophytes. In other words, when genes exhibited a moderate to high degree of gametophyte-biased expression, this was predominantly due to strong downregulation (silencing) of these genes in the sporophyte generation (Figure 1B).

Interestingly, in the Ectocarpales species more than 80% of the sporophyte-biased genes exhibited fold changes of between 2 and 6, whereas in the Laminariales species (which have a dominant sporophyte generation) a greater proportion of the sporophyte-biased genes exhibited very high fold changes between generations, with between 13% and 23% in *S. japonica* and *M. pyrifera,* respectively, exhibiting fold changes of more than 20 (Figure 1B; Table S2). Nevertheless, in all four species the majority of the generation-biased genes with very strong bias (FC>20) were gametophyte-biased (Figure 1B; Table S2), with as many as 50% of the gametophyte-biased genes in *S. lomentaria* belonging to this group (15% in *Ectocarpus* sp., 28% in *M. pyrifera* and 44% in *S. japonica*).

We also noted that, on average, sporophyte-biased genes were expressed at significantly higher levels than gametophyte-biased genes in all four species (Wilcoxon test, p<2e−12) (Figure 1C). In general, the complexity of each life cycle generation, both in terms of number of cell types and size of the organism, tended to be correlated with the number of generation-biased genes (Table S3). For instance, *S. lomentaria*, which has a dominant gametophyte generation, possessed the highest proportion of gametophyte-biased genes. Conversely, *M. pyrifera* and *S. japonica*, whose sporophytes are much larger than the gametophytes (Figure S1, Table S3), exhibit the highest proportion of sporophyte-biased genes.

### High turnover of generation-biased gene sets in the brown algae

Comparisons of orthogroups (OGs) containing generation-biased genes showed that they were poorly conserved between pairs of species (Figure 2A). Only 23% and 20% of the orthogroups (OGs) containing generation-biased genes were shared by the Ectocarpales and Laminariales species pairs respectively, and conservation between pairs of species from different orders was even lower (14% to 17%). Only 2% of the generation-biased OGs were conserved across all four of the study species (Figure 2A). Importantly, a large proportion of the generation-biased genes (between 36% and 63%) did not have orthologs in the genome of any of the other three study species nor in the genomes of four other distant Stramenopile species (Figure 2B; Table S4). We refer to these taxonomically-restricted genes hereinafter as “orphan” genes. The orphan genes were not included in the OGs analysis described above (Figure 2A) because most orphans are not members of an OG. The analysis of OGs therefore actually overestimated the degree to which generation-biased gene sets were conserved across species.

**Figure 2.**
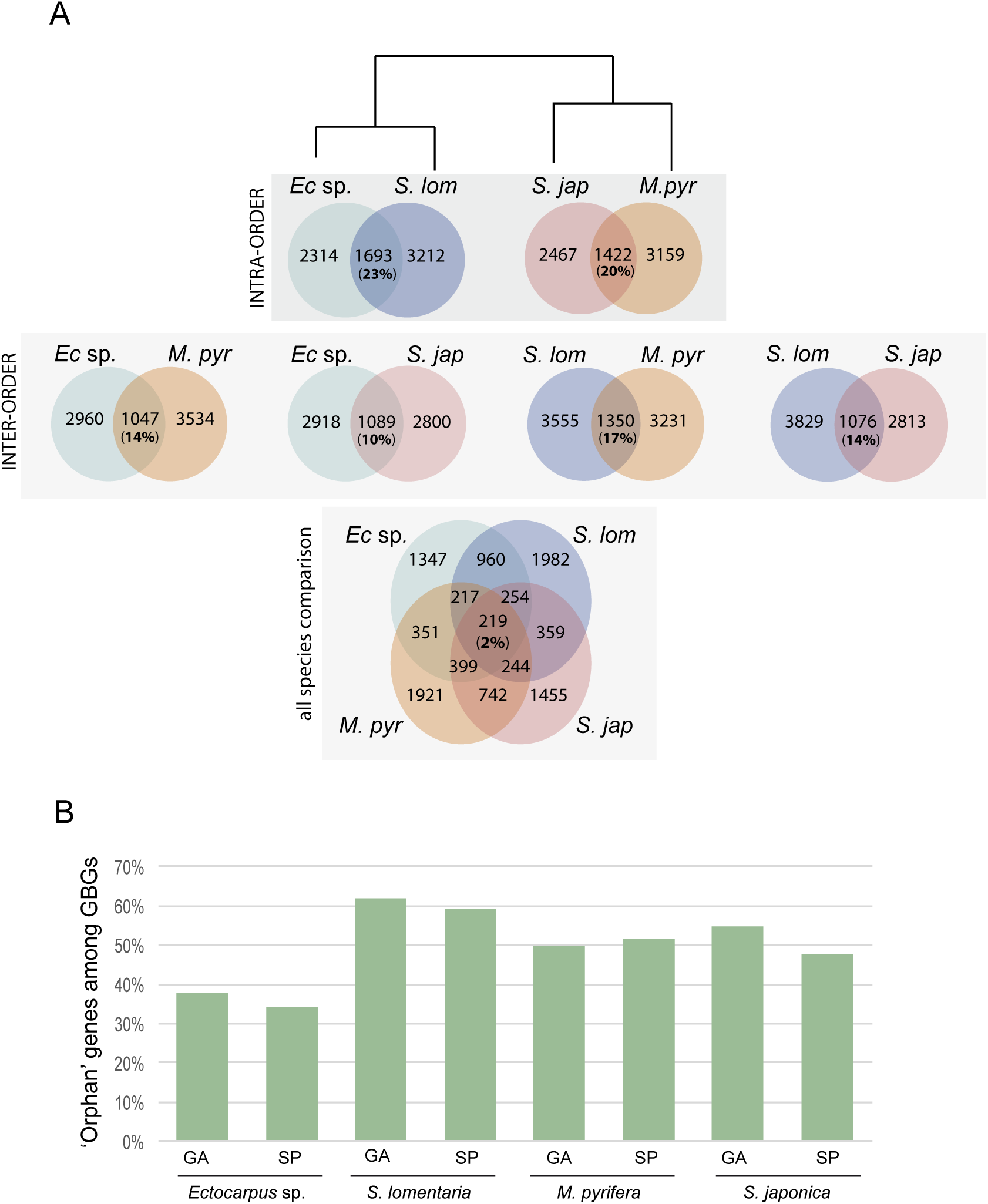
OGs with generation-biased genes are poorly conserved across brown algal species and the generation-biased gene sets include many orphan genes. **(A)** Shared OGs with generation-biased genes across the four-studied species. Venn diagrams representing the number of shared versus species-specific generation-biased OGs. Comparisons were made at several evolutionary distances. **(B)** Proportion of orphan (taxonomically-restricted) genes within the generation-biased gene sets of each of the four-studied species. GBGs: generation-biased genes.

The sets of gametophyte-biased genes in *Ectocarpus* sp., *S. lomentaria* and *S. japonica* and the set of sporophyte-biased genes in *S. japonica* were significantly enriched in orphan genes compared to the whole genome (Fisher test, p<0.03, p=0.0008, respectively), suggesting that *de novo* evolution has played an important role in the emergence of the generation-biased gene sets.

### Evolution of generation-biased expression of orthologous genes across brown algal species

To further analyse the evolutionary history of the generation-biased genes, we focused on genes for which there was a clear orthologous relationship across the four species. A large set of 15,888 orthologous groups (OGs), identified using OrthoMCL, was screened to identify 6,035 single copy orthologous genes with either 1:1:1:1 or 1:1:1:0 occurrence across the four brown algal species (see methods for details). We will refer to this set of OGs as “all single orthologues” (ASOs).

The ASO dataset was used to assess the conservation of generation-biased gene expression across the four species. Of the 6,035 ASOs, 4,489 (74%) included genes that were generation-biased in at least one of the species. However, only 35 gametophyte-biased genes and three sporophyte-biased genes consistently exhibited patterns of generation-biased expression across all four species (Figure 3A). The number of genes with conserved generation-biased expression increased to 175 gametophyte-biased and 81 sporophyte-biased when we took into account orthologous genes with generation-bias in three species (with the ortholog missing or unbiased in the fourth species) (Figure 3A). Fifteen percent of the ASOs (904 of the 6,035) exhibited discordant generation-biased expression patterns, so that, for example, the orthologue of a gene that was sporophyte-biased in one species was gametophyte-biased in at least one of the other three species (Figure 3A).

**Figure 3.**
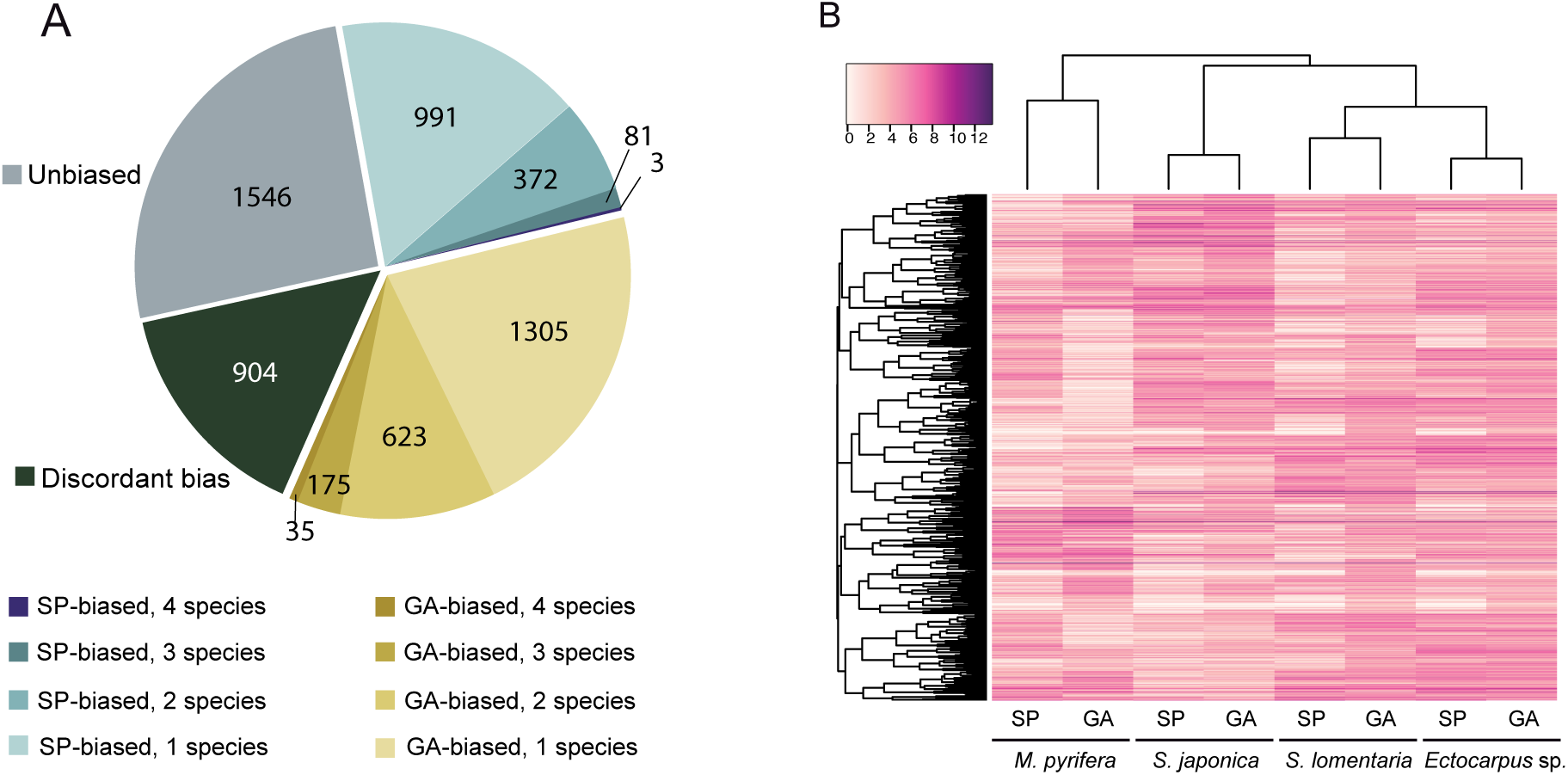
Conservation of generation-biased gene expression across species. **(A)** Numbers of ASOs showing unbiased, discordant bias or different degrees of shared bias between the four studied species. GA: gametophyte; SP: sporophyte **(B)** Hierarchical clustering and heatmap of gene expression for all the members of the 1:1:1:1 ortholog dataset with at least one generation-biased member in one of the studied species (Heatmap3 package, R). The dendogram was constructed using hierarchical clustering with 1000 bootstraps (pvclust package, R).

We used hierarchical clustering of expression levels for all the members of the 1:1:1:1 ASO dataset with at least one generation-biased member in one of the studied species to visualize global transcription patterns within and among the four species. In this analysis, samples clustered primarily by phylogenetic relatedness and not according to life cycle stage (Figure 3B), reflecting the low level of conservation of generation-biased expression patterns of gene expression across the lineages.

Taken together, these analyses indicated that the gametophyte-biased transcriptome tended to be more conserved than the sporophyte-biased transcriptome, but overall, generation-biased expression of the ASO dataset was extremely poorly conserved across the four brown algal species.

### Generation-biased gene expression within the Ectocarpales and Laminariales

To analyse divergences of generation-biased expression patterns within orders, we used the OrthoMCL analysis to identify single copy (1:1) orthologues shared either by the two Ectocarpales (6,438 OGs) or by the two Laminariales (5,061 OGs) species. These sets of 1:1 OGs were termed "pairwise single orthologues" (PSOs) (Table S5).

Between 22-34% (Ectocarpales) and 24-33% (Laminariales) of the generation-biased genes were PSOs, whereas the remaining generation-biased genes had no ortholog in the other species from the same order (Figure 4A). Between 11-22% (for both Ectocarpales and Laminariales) of the generation-biased genes gained either sporophyte- or gametophyte-biased expression in one of the species (Figure 4A), whereas discordant generation-biased expression was observed for 1.5-5% (Ectocarpales) and 2-5% (Laminariales) of the generation-biased genes (Figure 4A).

**Figure 4.**
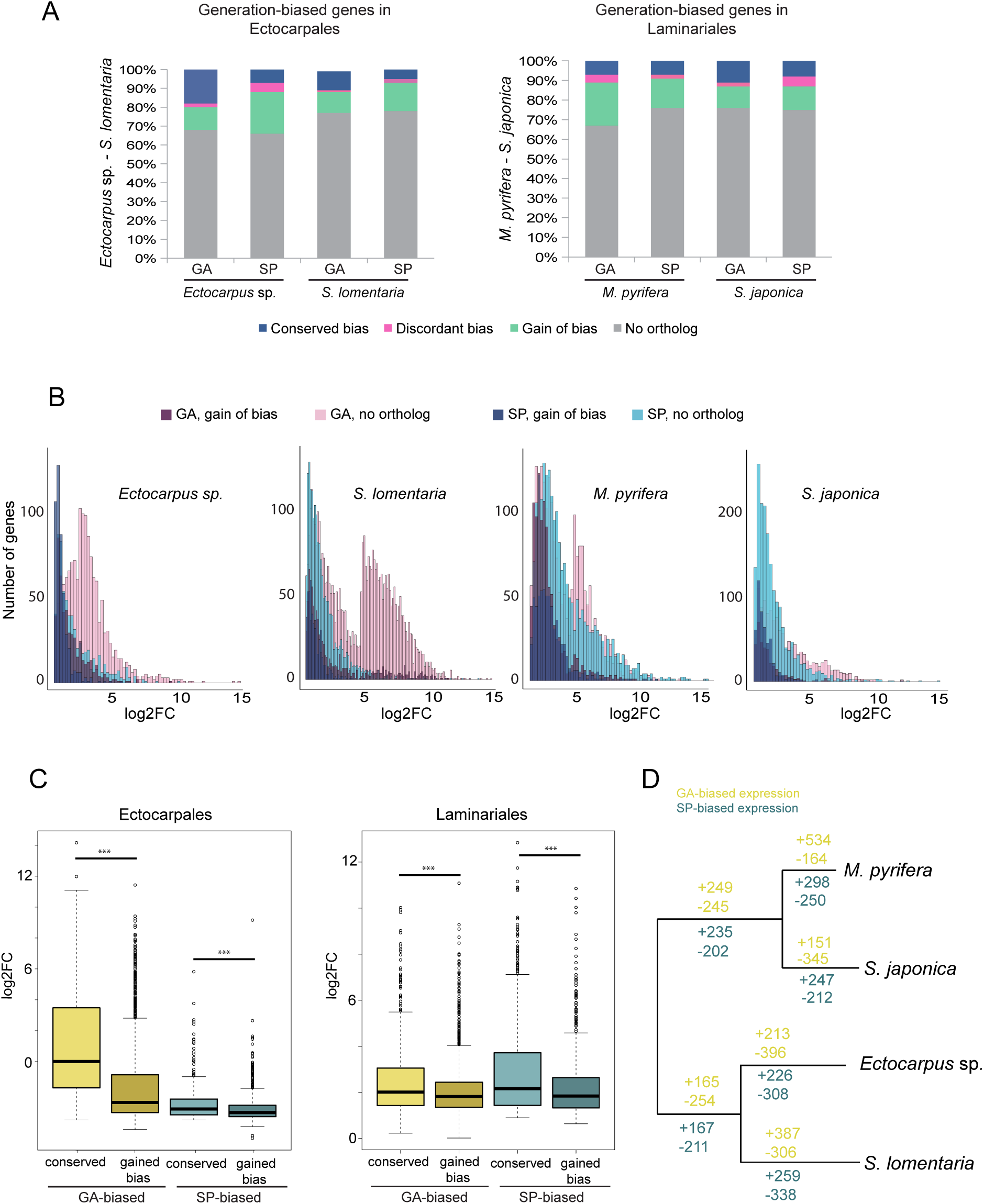
Conservation of generation-bias across the Ectocarpales (Ectocarpus sp. and S. lomentaria) and Laminariales (S. japonica and M. pyrifera). **(A)** Proportion of the PSOs among generation-biased genes. In grey, genes which had no ortholog in the other species of the same order, colour indicates PSOs. For the PSOs, genes with ‘Conserved bias’ (dark blue) exhibited the same bias in the two species. Genes with ‘Discordant bias’ (pink) were gametophyte-biased in one species and sporophyte-biased in the other species. ‘Gained bias’ genes (green) were generation-biased in one species but not in the other species. **(B)** Distribution of magnitude of generation-bias (log2 fold changes) for PSOs that gained bias (ortholog is unbiased in the other species) and species-specific genes with biased expression (no ortholog in the other species). Orthology was established per lineage in pairwise comparisons between Ectocarpus sp.-S. lomentaria and S. japonica-M. pyrifera. **(C)** Overall level of generation-biased expression (log2FC) for PSOs that are conserved versus PSOs that gained bias in Ectocarpales and Laminariales. **(D)** Representations of gene gain/loss events across the branches of the Ectocarpales and Laminariales phylogeny. Expected numbers of events are based on multiple stochastic mappings (see methods for detail).

Overall, species-specific gametophyte-biased genes presented greater bias, measured as fold change (Figure 4B). This was particularly pronounced in *S. lomentaria,* where about half of the species-specific genes with gametophyte-bias had expression levels at least 50 times higher (log_2_FC>5.7) than in the sporophyte. In Laminariales, conversely, it was the species-specific genes with sporophyte-biased expression that presented overall higher fold changes (Figure 4B). In other words, newly emerged, species-specific genes showed a strong magnitude of bias in gametophytes of Ectocarpales and sporophytes of Laminariales.

A correlation was observed between the level of bias and the conservation of generation-biased genes within each lineage (Ectocarpales and Laminariales), i.e. the mean fold change of genes that were conservatively generation-biased (both gametophyte-biased or both sporophyte-biased) across the species pairs was significantly higher than that of genes were generation-biased in one of the species but unbiased in the other (Figure 4C; Wilcoxon test, p<1e-05). Taken together, these analyses suggested that the rapid evolution of generation-biased gene sets involves not only the emergence of new generation-biased genes but also the emergence, in species-specific fashion, of novel generation-biased expression patterns associated with existing orthologous genes.

### Evolutionary history of generation-biased gene sets

We used a phylogenetic stochastic mapping approach to investigate the evolution of generation-biased gene expression. Phylogenetic stochastic mapping allows reconstruction of the history of trait changes (in our case, generation-bias) based on the estimation of the probabilities and expectations of gain and loss events for each branch of an underlying phylogenetic tree [23]. Rates of gain and loss were equal for both gametophyte- and sporophyte-bias, as determined by a likelihood ratio test between the ER and ARD models (all p-values >0.05). The stochastic mapping results highlighted widespread and rapid turnover of generation-biased genes during evolution of the Laminariales and Ectocarpales (Figure 4D). Specifically, more events of gain of bias were observed for gametophyte-biased genes, and, conversely, sporophyte-biased genes presented more events of loss of bias. Overall, gametophyte-biased genes presented a more rapid turnover with a higher total number of events compared with sporophyte-biased (3409 versus 2953).

### Duplicated generation-biased genes

Gene duplication and generation-specific co-option of paralogs may be a mechanism to resolve potential generation antagonism due to evolutionary divergence between the two generations. Analysis of in-paralogs identified by OrthoMCL indicated that the generation-biased gene sets of *Ectocarpus* sp., *S. lomentaria* and *M. pyrifera* were not enriched in members of duplicated gene pairs compared with the rest of the genes in each genome (Fisher test, p=1). However, duplicated genes constituted 32 and 36% of gametophyte and sporophyte-biased genes in *S. japonica* (Table S4, Figure 5), which was significantly more than expected by chance (Fisher test, p<2e-16 for both comparisons).

**Figure 5.**
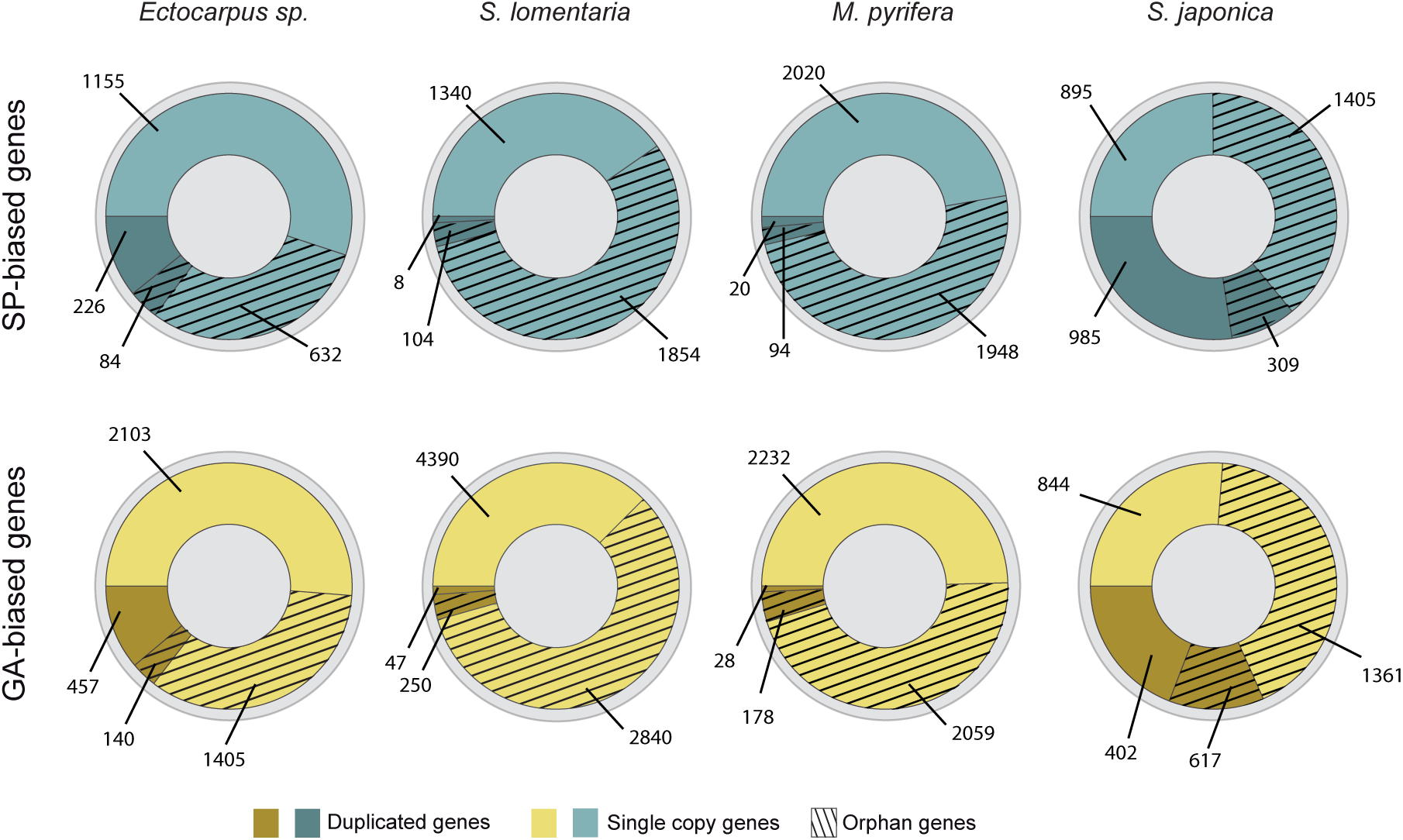
Proportion of single copy versus duplicated genes and amount of orphan (taxonomically restricted) genes among gametophyte and sporophyte-biased genes in each of the brown algal species.

Discordant bias was observed for 10-44% of generation-biased in-paralog pairs. *S. japonica* had significantly more generation-biased in-paralog pairs that showed discordant bias than the other three species (Chisqr test, p<0.007). The set of in-paralogs with discordant bias was completely different for each species, indicating that duplication of genes followed by acquisition of two, opposite generation-biased expression patterns by the resulting in-paralogs occurred independently in each of the four lineages.

### Predicted functions of generation-biased genes

An analysis of gene ontology (GO) terms associated with the generation-biased genes was carried out using Blast2GO [24] to search for enrichment in particular functional groups. First, Blast2GO analysis was carried out for each species, in order to relate gene function to the phenotypic generation dimorphisms specific to each species.

The GO terms associated with *M. pyrifera* and *S. japonica* sporophyte-biased genes were enriched in biological processes related to reproduction, carbohydrate metabolism, protein modification, growth and development, signalling, cell communication, response to external stimulus and homeostasis (Fisher exact test, p<0.05, Table S6). Interestingly, a similar set of GO terms was enriched for the gametophyte-biased genes of *S. lomentaria*, in addition to categories related to sexual reproduction and cilium motility (Figure S2, Table S6). This result suggest that similar genetic processes are at work in the morphologically complex, long-lived, dominant generations of these three species, despite the large morphological differences between gametophytes of *S. lomentaria* and sporophytes of Laminariales and the limited number of shared generation-biased genes.

Analysis of the gametophyte-biased genes from all the studied species identified six GO terms related to biological processes that were consistently significantly enriched in all species (Fisher exact test p<0.05). These terms included microtubule and flagellar movement-related categories and corresponded to between 10% and 50% of gametophyte-biased genes with assigned ontology (Table S7). Three GO terms were consistently enriched for the sporophyte-biased genes of all the studied species. These terms, which were related to carbohydrate metabolism and small GTPase signalling processes, corresponded to between 10% and 20% of the sporophyte-biased genes in each species.

### Structural characteristics of the generation-biased genes

Several structural characteristics (GC and GC3 content, coding region size and intron number) were compared between sporophyte-biased, gametophyte-biased and unbiased genes (Figure S3; Table S8). Gametophyte-biased genes tended to have longer coding regions, to possess more introns and to have a lower GC3 content than unbiased genes in all four species (Wilcoxon test, p-value<6e-04). In contrast, sporophyte-biased genes did not present a consistent trend in relation to the unbiased genes in any of the species studied. Excluding orphan genes from the analysis did not significantly change the results indicating that the structural characteristics of gametophyte-biased genes were not due to the fact that these genes are evolutionary younger (Figure S4).

### Evolutionary features of the generation-biased genes

The evolutionary dynamics of generation-biased genes was investigated by calculating ratios of pairwise synonymous to nonsynonymous substitution rates and comparing these data with gene expression divergence (see Methods). This analysis, which was applied to all the PSOs that could be examined for each order (Ectocarpales and Laminariales), indicated that gametophyte-biased genes evolve faster (i. e., had higher dN/dS ratios) than unbiased or sporophyte-biased genes in both Ectocarpales species (Figure 6A; Wilcoxon test, p<1e-11) and in *M. pyrifera* (Wilcoxon test p<0.005). The kelp *S. japonica* was the only exception, no significant difference was observed between the rates of evolution of gametophyte-biased and unbiased genes for this species (Figure 6A).

**Figure 6.**
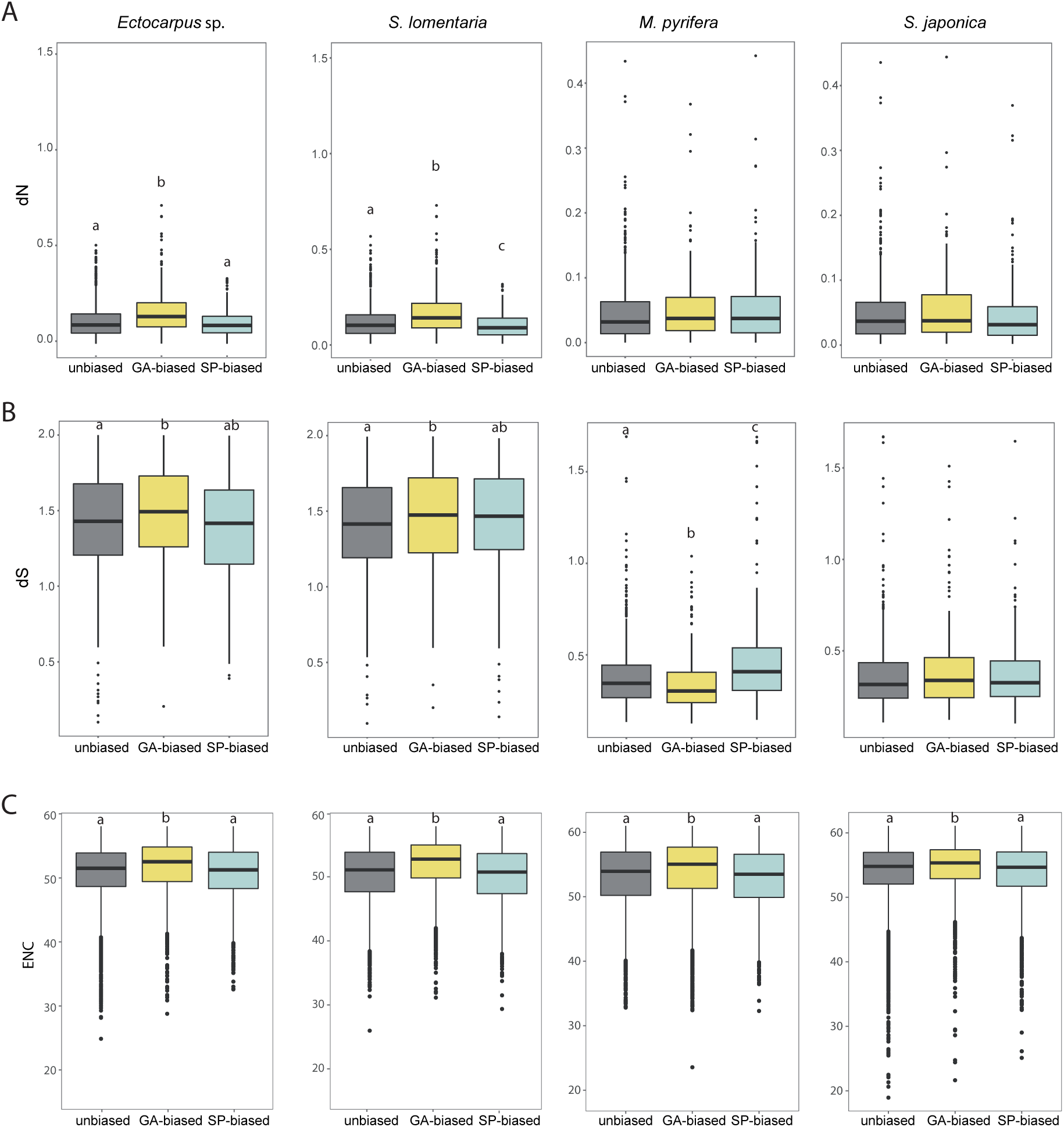
Evolution of generation-biased genes. (A) Evolutionary rates measured as dN/dS between species pairs (Ectocarpus sp./S. lomentaria, M. pyrifera/S. latissima) for unbiased, gametophyte-biased and sporophyte-biased genes in the four brown algal species. (B) Gene expression divergence measured as Euclidean distances for unbiased, gametophyte- and sporophyte-biased genes in each of the four brown algal species. Different letters above the plots indicate significant differences (Wilcoxon test; P<0.05).

Sex-biased genes have been shown to exhibit accelerated rates of evolution in *Ectocarpus* sp. [25] and many of the gametophyte-biased genes also exhibit a sex-biased pattern of expression (809 genes or about 20%) because this is the sexual generation of the life cycle. However, when these sex-biased genes were removed from the dN/dS analysis, the gametophyte-biased genes still showed faster evolutionary rates than unbiased or sporophyte-biased genes (Figure S5A; Wilcoxon test p-value=6.3e-13 for the gametophyte versus unbiased comparison, p-value=2e-09 for the gametophyte versus sporophyte-biased comparison) indicating that the faster evolutionary rates of gametophyte-biased genes were not solely due to the presence of sex-biased genes. Similar results were obtained when we subdivided the generation-biased genes into conserved bias and species-specific bias (i.e. genes that showed generation bias in one species but were unbiased in the other) (Figure S5B, S5C). This latter analysis suggested that the faster evolutionary rates of gametophyte-biased genes were not correlated with the degree of conservation of expression across the lineages.

The higher evolutionary rates of gametophyte-biased genes were due to the accumulation of both synonymous and non-synonymous changes (Figure S6A, S6B). There was no correlation between dN/dS and fold change of generation-biased gene expression between generations (Spearman’s rho=-0.037). Codon usage bias (CUB), measured as the effective number of codons (ENC), indicated that gametophyte-biased genes had significantly lower codon usage bias (i.e. higher ENC) than sporophyte-biased and unbiased genes in both Ectocarpales and Laminariales (pairwise Wilcoxon test p<3.0e-11) (Figure S6C). We found no significant difference in CUB between sporophyte-biased and unbiased genes.

Overall, order-specific genes (i.e., ‘young’ genes) exhibited higher evolutionary rates (dN/dS) than genes conserved across the four species (‘old’ genes) for both the Ectocarpales (Wilcoxon test, p-value=2.8e-13) and Laminariales (Wilcoxon test, p-value=6.5-06), suggesting that younger genes are evolving more rapidly (Figure S7).

To assess whether increased protein divergence rates were due to increased positive selection or relaxed purifying selection, we performed a maximum likelihood analysis using codeml in PAML4. In addition to the four study species, we searched for orthologs of the ASOs in published transcriptome data for three additional *Ectocarpus* species (*E. fasciculatus,* un unnamed *Ectocarpus* species from New Zealand and *Ectocarpus siliculosus)* and the recently published genome of *C. okamuranus* [26]. The 522 conserved orthologs identified by this analysis included 393 orthologs that exhibited generation bias in at least one of the studied species. For 42 of these comparisons both pairs of models (M1a-M2a, M7-M8) suggested positive selection and a total of 111 genes were predicted to be evolving under positive selection as indicated by the model M7-M8 alone (Table S9). Among the 111 genes identified by the M7-M8 model, 83 exhibited generation-bias in at least one species. Taken together, our analysis is consistent with the idea that a subset of the generation-biased genes exhibit signatures of positive selection, although the set of generation-biased genes was not significantly enriched in genes that were predicted to be under positive selection (Fisher’s exact test, P=0.8045).

When patterns of gene expression were considered, measured as Euclidian distances for PSOs within each order, sporophyte-biased genes showed overall significantly larger divergence that unbiased or gametophyte-biased genes (Figure 6B; Wilcoxon test p<3e-7). Furthermore, gametophyte-biased genes exhibited the most conserved expression patterns within each order, even in comparison to unbiased genes.

Taken together, our data suggests that gametophyte- and sporophyte-biased genes have distinct patterns of evolution: gametophyte-biased genes tend to exhibit rapid evolution of their coding sequence whereas sporophyte-biased genes tend to exhibit changes to their patterns of expression.

## DISCUSSION

Here, we have used four phylogenetically-diverse brown algal species with different levels of generation dimorphism and complexity to investigate genome-wide generation-biased gene expression patterns and to assess the potential role of generation-specific selection in shaping these patterns of gene expression.

### Differential gene expression underlies phenotypic dimorphism between life cycle generations

Between 35% and 46% of the genome of the brown algae species studied here was differentially regulated during the gametophyte and sporophyte generations of the life cycle. This is substantially more than in the moss *Funaria hygrometrica,* where only 4% of the genome is differentially expressed [27], and even exceeds the situation in *Arabidopsis,* where about 30% of the genome is generation-biased [28]. Comparative analysis between these two land plants showed that the relative proportion of generation-biased genes assigned to the two life cycle generations was lower in the moss than in *Arabidopsis* [27], consistent with the lower level of phenotypic dimorphism between generations in the former. Likewise, we found that for the brown algae studied here the relative numbers of generation-biased genes during each generation was broadly correlated with size differences between the two generations. Despite this tendency, however, the absolute number of gametophyte-biased genes was consistently greater than the number of sporophyte-biased genes across all four studied species, even in kelps where the gametophyte is much less complex, morphologically, than the sporophyte. This tendency is difficult to explain but we note that a previous analysis using *Ectocarpus* sp. also indicated that the gametophyte developmental program deployed more genes than the sporophyte program [4]. Based on this difference it was proposed that the switch to the sporophyte program involves predominantly gene repression. Again, this hypothesis is consistent with our analysis indicating that gametophyte-biased gene expression tends to be the result of downregulation during the sporophyte generation.

### Generation-conflict and generation-biased gene expression

In many brown algal species, free-living gametophytes and sporophytes display extensive morphological and physiological dimorphisms (reviewed in [1,29,30]) and this phenotypic diversity reflects in many cases different ecological niche preferences for gametophytes and sporophytes (e.g.[14,31]). Gametophyte and sporophyte development and function are controlled by a common genome, with a large number of genes carrying out functions during both generations. When there are marked morphological and physiological differences between the two generations, as is the case for most of the species studied here, this can lead to conflict due to genes being submitted to different selection pressures during the different generations of the life cycle. Generation-biased gene expression is one mechanism to reduce inter-generational conflict, allowing gene products to be targeted specifically to one generation (although this does not necessarily mean that every generation-biased gene arose due to generation antagonism). Gene duplication, followed by acquisition of generation-bias through neo-functionalization, can play an important role in the resolution of generation conflict and we found some evidence for this, at least for one of the kelp species. Note that gene duplication followed by neo-functionalisation has also been proposed as one of the mechanisms that allow resolution of sexual antagonism (reviewed in [17]).

### Generation-biased genes turnover rapidly during the evolution of the brown algae

Perhaps the most striking result of our analysis is the remarkably limited number of generation-biased genes that were shared by all the four of the studied species, indicating a rapid turnover of life cycle-biased genes in brown algae. This turnover appears to be due to a combination of two processes: emergence of new genes with generation-bias and gain/loss of bias in existing, orthologous genes. The former mechanism appears to have been of paramount importance as between 36% and 63% of generation-biased genes were orphans, depending on the species. Phylogenetic stochastic mapping results were consistent with rapid loss and gain of bias in orthologous genes, and highlighted a particularly high rate of turnover for gametophyte-biased genes.

The ancestor of brown algae is thought to have alternated between multicellular, isomorphic gametophyte and sporophyte generations without a clearly dominant generation [22]. From this morphologically simple ancestor, there would have been a tendency, in most brown algal lineages, to evolve towards increased complexity of either the gametophyte or the sporophyte generation [22]. Our data indicate that this increase in size and developmental complexity was accompanied by an overall increase in the proportion of the transcriptome that become gametophyte- or sporophyte-biased, depending on the lineage.

Recent analysis of *Ectocarpus* sp. developmental mutants have indicated that the evolution of the sporophyte and gametophyte genetic programs involved both co-option of genetic programs from one generation to the other and generation-specific innovations [34,35]. We observed that a subset of the expressed genes in the four brown algal species exhibited switching of bias between life cycle generations in the different lineages, in line with the idea that the evolution of the generation-specific developmental programs in the brown algae has, to some extent, involved sharing of genes between generations. But importantly, our evidence indicates that the evolution of brown algal gametophyte and sporophyte developmental programs has been predominantly driven by the emergence of lineage-specific, orphan genes, suggesting a more important role for generation-specific innovations during the evolution of the sporophyte and gametophyte genetic programs.

The evolutionary origins of the sporophyte and gametophyte developmental programs in land plants have been intensively studied, particularly with regard to the question of whether each generation has independently evolved its own developmental pathways or, alternatively, whether there has been recruitment of developmental programs from one generation to the other during evolution [36–38]. It is currently thought that the developmental networks that implement land plant sporophyte programs were mainly recruited from the gametophyte generation, which was initially the dominant generation [27,36,39] although there is also evidence that there have been sporophyte-specific innovations [27,40]. We suggest however, based on the observations presented here for the brown algae, that it may be an oversimplification to think in terms of one generation gradually recruiting programs from the other generation, and that a more extensive sampling of generation-biased gene sets in land plants may reveal a more dynamic situation involving important amounts of both lineage-specific gene evolution and lineage-specific switching of generation-biased expression patterns.

Interestingly, despite the marked differences between the generation-biased gene sets of the four studied brown algae, the enriched GO terms associated with genes expressed during the more morphologically complex, long-lived, dominant generation tended to be similar. The predicted functions of both the sporophyte-biased genes of kelps and the gametophyte-biased genes of *S. lomentaria* were enriched in GO terms associated with polysaccharide and cell wall biosynthesis, developmental processes, cell signalling and cell communication. These conserved, enriched GO terms could reflect developmental and morphological processes common to dominant life cycle generations, such as extended multicellular growth.

### Rapid evolution of the coding regions of gametophyte-biased genes

On average, gametophyte-biased genes were found to be evolving significantly more rapidly (higher dN/dS) than sporophyte-biased and unbiased genes in all the species studied except *S. japonica*. This was surprising because purifying selection is expected to be more efficient for genes expressed during the haploid phase of the life cycle due to the absence of masking of recessive and partially recessive mutations [42,43]. However, accelerated evolution of gametophyte-biased genes has also been previously reported in land plant systems [28,41] and a number of hypothesis have been put forward to explain this phenomenon. It has been suggested, for example, that gametophyte-biased genes are under relaxed constrain because of lower expression breadth and low level of tissue complexity [27,41,44–46]. This hypothesis is unlikely to explain our observations because in *S. lomentaria* the gametophyte generation is the dominant phase of the life cycle and is larger and more complex than the sporophyte. It has also been proposed that strong selection on reproductive traits during gametogenesis (i.e., during the gametophyte-generation) may explain the faster rates of evolution of gametophyte-biased genes in plants [41]. However, when sex-biased genes were excluded from the *Ectocarpus* sp. gametophyte-biased gene dataset, the remaining genes still exhibited a significantly higher rate of evolution than that of unbiased or sporophyte-biased rates. Finally, land plant gametophyte-biased gene sets are enriched in young genes [34] and this may affect evolution rate as young genes are known to evolve more rapidly. However, rapid evolution of young genes is unlikely to explain the faster evolutionary rates of brown algal gametophyte-biased genes because the gametophyte- and sporophyte-biased gene sets contained similar numbers of young (faster evolving) genes.

We did note, however, that gametophyte-biased genes have overall lower levels of expression than sporophyte-biased genes, and expression levels have been negatively correlated with evolutionary rates [47–49]. Moreover, since that gametophyte-biased genes turn over more rapidly than sporophyte-biased genes, one interesting possibility is that gametophyte-biased genes may be less associated with complex gene interaction networks, and therefore be more dispensable and thus under less constraint [50–52]. More information about gene interaction networks will be needed for the brown algae in order to test this hypothesis.

### Sporophyte-biased and gametophyte-biased genes exhibit different patterns of evolution

In contrast to the gametophyte-biased genes, sporophyte-biased genes did not exhibit overall accelerated rates of evolution of their coding sequences but they did exhibit significantly higher levels of diversification of expression patterns (measured as Euclidean distance), compared to both unbiased and gametophyte-biased genes. Therefore, whilst the gametophyte-biased genes exhibited accelerated evolution of their coding regions, the sporophyte-biased genes appeared to have experienced accelerated evolution of their regulatory sequences. Decoupling of protein sequence evolution and expression pattern evolution has been observed in other eukaryotes (e.g. [50,51], but see [52,53]) but it is not clear why the mechanisms of evolution should differ between gametophyte-biased and sporophyte-biased genes in these brown algal species. It has been suggested that mutations that change protein sequences and mutations affecting gene regulation play different roles during evolution, with genes involved in physiological traits tending to exhibit the former and genes involved in morphological traits evolve primarily in terms of gene expression [58,61]. An in-depth functional analysis of brown algae genes using experimental approaches will be crucial to understand if the different evolutionary modes of gametophyte- and sporophyte-biased genes are associated with different functions of the gene networks underlying each generation.

## CONCLUSIONS

This study afforded the first comparative analysis of generation-biased gene expression across several species with complex life cycles to understand the role of generation-specific selection in shaping patterns of gene expression and divergence. Our analyses revealed that an extensive proportion of the genome exhibits generation-biased expression in the brown algae and the relative proportion of genes that are generation-biased was correlated with the degree of phenotypic dimorphism between generations. Life cycle-biased genes turn over very rapidly during evolution due to a combination of two processes: the gain/loss of generation-biased expression by orthologous loci and the emergence, *de novo,* of genes with generation-biased expression. Our results are consistent with the idea that the evolution of the genetic program associated with each generation appears to have involved the recruitment of genes across generations but the exceptionally high number of generation-biased orphan genes emphasizes an important role for generation-specific developmental innovations in each lineage. Finally, our analysis has revealed that gametophyte and sporophyte exhibit overall distinct evolutionary modes, with gametophyte-biased genes evolving rapidly predominantly at the level of their sequence and sporophyte-biased genes diverging mostly at the level of their patterns of expression.

## METHODS

### Biological material and generation of genomic and transcriptomic sequence data

The algal strains used, sequencing statistics and accession numbers are listed in Table S10. We used published RNA-seq datasets for gametophytes and sporophytes of the model brown alga *Ectocarpus* sp. [25,62], *S. japonica* (Teng et al., 2017), *M. pyrifera* (Lipinska et al., 2017) and for gametophytes of *S. lomentaria* (Lipinska et al., 2015). *S. lomentaria* sporophytes (strain Zy2) were derived from a controlled laboratory cross between the Asari6 female gametophyte and the Asari9 male gametophyte. Both gametophytes were field collected. Sporophyte clones were grown in 20°C 14h:10h light:dark conditions, in half-strength Provasoli enriched seawater [63] which allowed them to be maintained as immature thalli (absence of meiotic structures). For the gametophyte and sporophyte of *Ectocarpus* sp., gametophyte and sporophyte of *S. lomentaria* and gametophyte of both kelps we used whole thallus for RNA extractions. Total RNA was extracted using the Qiagen Mini kit (http://www.qiagen.com) as previously described (Coelho, et al. 2012). RNA was sequenced with Illumina HiSeq 2000 paired-end technology with a read length of 125 bp (Fasteris, Switzerland) and is available under the accession number detailed in Table S10.

For the sporophytes of kelps, we used RNA-seq data produced from replicate samples of blades tissue of adult individuals from natural populations (SRA references provided in Table S10).

Sets of reference genes for each species were derived from the published genomes of *Ectocarpus* sp. and *S. japonica* [64,65] or from draft genome assemblies for *S. lomentaria* and *M. pyrifera* [62] with gene prediction based on mapping of RNA-seq data using Stringtie [66]. Quality filtering of the raw reads was performed with FastQC (http://www.bioinformatics.babraham.ac.uk/projects/fastqc), and adapter sequences were trimmed using Trimmomatic (leading and trailing bases with quality below 3 and the first 12 bases were removed, minimum read length 50 bp [67]. *Ectocarpus sp.* and *S. japonica* reads were aligned to the reference genomes [65,68] using Tophat2 [59–61]. Gene models for *S. lomentaria* and *M. pyrifera* were predicted using Transdecoder (http://transdecoder.sf.net) based on mapping of the RNA-seq data (Tophat2) to the draft genome sequences generated previously for these two species [62]. Gene expression levels were represented as TPMs. Genes with expression values below the 5^th^ percentile of all TPM values calculated per species were considered not to be expressed and were removed from the analysis.

### Identification of generation-biased and generation-limited genes

The filtering steps described above yielded a set of expressed genes in the transcriptome that were then classified based on their generation-expression patterns. Genes were considered to be gametophyte-biased or sporophyte-biased if they exhibited at least a 2-fold difference in expression between generations with a false discovery rate (FDR) of < 0.05. Generation-biased genes were defined as generation-limited when the TPM was below the 5^th^ percentile for one of the generations.

### Gene orthology

OrthoMCL (Fischer, 2011) was used to assess orthologous relationships between the genes of the four-studied species (blastp, e-value < 1e-5). OrthoMCL identified a total of 15,888 orthogroups (OGs), of which 3,290 contained only one gene per species and therefore represented the set of 1:1:1:1 OGs. An additional 2,745 OGs had only one member in three of the studied species but no ortholog (i.e. the gene was missing) in the fourth species (1:1:1:0 OGs). We considered that these 1:1:1:0 OGs, which most likely represent single copy ancestral genes that were lost in one of the species, also provided useful information about conservation of generation-biased gene expression because they consisted of members from two different orders (Ectocarpales and Laminariales). We therefore combined the two sets of OGs (1:1:1:1 and 1:1:1:0) to create the “all single orthologues” (ASO) dataset, which was composed of a total of 6,035 OGs. The ASO dataset was employed to assess conservation of generation-biased gene expression across the four-studied species.

For pairwise comparisons within orders, we selected OGs that contained only one member in each of the two species (6,438 OGs for the Ectocarpales and 5,061 OGs for the Laminariales). We refer to the OGs in these datasets as “pairwise single orthologues” (PSOs).

Orphan (or *de novo*) genes (i. e., taxonomically restricted genes) were defined as genes present in the genome of only one species and having no BLASTp match (10-4E value cutoff) with a range of other stramenopile genome-wide proteomes from public databases (indicating that they are likely to have evolved since the split from the most recent common ancestor): the brown alga *Cladosiphon okamuranus* [26] the eustigmatophyte *Nannochloropsis gaditana* [72], the pelagophyte *Aureococcus anophagefferens* [64] and the diatom *Thalassiosira pseudonana* [74]. Duplicated genes were identified in each species using the in-paralog list generated by OrthoMCL.

### Prediction of gene function

InterProScan [75] and BLAST2GO [24] were used to assign protein function annotations to genes in all four-studied species. Fisher’s exact test with a p*-*value cutoff of 0.05 was used to detect enrichment of specific GO-terms in various groups of generation-biased genes.

The visualization of gene ontology data used for Figure S2 was generated using Revigo [76].

### Evolutionary analysis

To estimate the evolutionary rates (non-synonymous to synonymous substitutions, dN/dS) for generation-biased and unbiased genes, pairwise analyses were carried on the PSOs for each order (Ectocarpales and Laminariales). Orthologous protein sequences were aligned with Tcoffee (M-Coffee mode [77], the alignments curated with Gblocks [78] and then translated back to nucleotide sequence using Pal2Nal [79] or TranslatorX [80]. Sequences that produced a gapless alignment exceeding 100 bp in length were retained for pairwise dN/dS (ω) analysis using Phylogenetic Analysis by Maximum Likelihood (PAML4, CodeML, F3×4 model of codon frequencies, runmode = −2) [81]. Genes with saturated synonymous substitution values (dS > 2) were excluded from the analysis.

The positive selection analysis was carried out using CodeML (PAML4, F3×4 model of codon frequencies) using additional orthologs of the 1:1 best ortholog set from OrthoMCL found in the transcriptomes of three *Ectocarpus* species (*E. fasciculatus,* an unnamed *Ectocarpus* species from New Zealand and *E. siliculosus;*) and in the genome of *Cladosiphon okamuranus* [26]. The analysis was therefore based on data from seven species in total. Protein alignment and curation was performed as described above. Gapless alignments longer than 100 bp containing sequences from at least three species were retained for subsequent analysis. CodeML paired nested site models (M1a, M2a; M7, M8) [81] of sequence evolution were used and the outputs compared using the likelihood ratio test. The second model in each pair (M2a and M8) is derived from the first by allowing variable dN/dS ratios between sites to be greater than 1, making it possible to detect positive selection at critical amino acid residues.

The effective number of codons (ENC) was calculated using ENCprime [82] with ribosomal genes as background nucleotide composition.

### Euclidean distances

Euclidean distances were estimated for all the PSOs for each of the two orders (Ectocarpales and Laminariales) following the approach of [83]. The following formula was used:

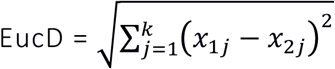

Where x_ij_ is the expression level of the gene under consideration (TPM) in species i (i.e. species 1 or species 2) during stage j (i.e. gametophyte or sporophyte) and k is the total number of stages (i.e. two, gametophyte and sporophyte). All statistical analysis was performed using RStudio (R version 3.4.2).

### Stochastic mapping approach to assess the evolutionary dynamics of generation-biased expression

We conducted an evolutionary analyses of the presence and absence of generation-biased gene expression as a dynamic between gain and loss of phyletic patterns [84]. To estimate the evolutionary dynamics of each event we tested whether the rates of gain (0→1) and loss (1→0) of bias were equal (ER model) or different (ARD model), and implemented stochastic mapping for each gene using the *phytools* R package [85]. The number of events on each branch only included those transitions that effectively produced a change of state at the start and end of the specified branch.

## DECLARATIONS

### Ethics approval and consent to participate

not applicable

### Availability of data and materials

All data generated or analysed during this study are included in this published article and its supplementary information files. Details of SRA references for sequence data are included in Table S10. All the datasets used and analysed during the current study are also available from the corresponding author on reasonable request.

### Author’s contributions

AL, AFP and KK prepared the biological material and performed experiments. AL and MS performed the computational analysis. AL, MS and SMC analysed data. SMC wrote the manuscript with valuable input from AL and JMC. SMC coordinated the study. All authors read and approved the final manuscript.

## Acknowledgements

The authors thank Nicolas Salemin (University of Lausanne) for fruitful discussions and advice on phylogenetic methods, Dieter Muller, Erasmo Macaya and Delphine Scornet for providing the photographs of *M. pyrifera* and *Ectocarpus* sp. shown in Figure S1.

## ADDITIONAL FILES

### Additional File 1 (.xlx): Supplemental tables

**Table S1.** Phenotypic differences between gametophytes (GA) and sporophytes (SP) of the four-studied species.

**Table S2.** Gametophyte- and sporophyte-biased genes identified for each of the four studied species (DEseq2 FC<2, padj<0.05, TPM>5th percentile).

**Table S3.** Generation-biased gene expression and morphological complexity of sporophytes and gametophytes across the four-studied species. GA: gametophyte; SP: sporophyte.

**Table S4.** Number of duplicated, single copy and orphan genes among the generation-biased genes in each of the studied brown algal species. SP: sporophyte; GA: gametophyte.

**Table S5.** Orthology statistics based on the OrthoMCL analysis. GBGs (generation-biased genes); GA (gametophyte); SP (sporophyte); PSOs (pairwise single orthologs); ASOs (all single orthologs).

**Table S6.** Enriched gene ontology (GO) terms associated with the generation-biased genes (GBGs) identified for each of the studied species (Fisher test p<0.05).

**Table S7.** Enrichment in GO terms common to all gametophytes and sporophytes.

**Table S8.** Structural characteristics of the generation-biased genes identified for each of the studied species

**Table S9.** PAML codeml analysis with the F3X4 substitution model. UB: unbiased; GA: gametophyte-biased, SP: sporophyte-biased.

**Table S10.** Strains used in this study and DNA and RNA sequencing data statistics.

### Additional file 2 (.doc): Supplemental Figures

**Figure S1.** Brown algal species used in this study. **(A)** *Ectocarpus* sp. sporophyte **(B)** *Ectocarpus* sp. gametophyte **(C)** *S. lomentaria* gametophyte; **(D)** *S. lomentaria* sporophyte; **(E)** *S. japonica* sporophyte; **(F)** *M. pyrifera* sporophyte; **(G)** *S. japonica* gametophyte; **(H)** *M. pyrifera* gametophyte.

**Figure S2.** Visualisation of GO terms associated with generation-biased genes for *S. lomentaria* gametophytes and *S. japonica* and *M. pyrifera* sporophytes. The scatterplot shows the cluster representatives (i.e. terms remaining after the redundancy reduction) in a two-dimensional space derived by applying multidimensional scaling to a matrix of the GO terms’ semantic similarities [76]. Bubble colour indicates p values and bubble size indicates the number of genes assigned with each GO term.

**Figure S3.** Structural characteristics (% GC, %GC3, CDS size and intron number) of unbiased, gametophyte- and sporophyte-biased genes across brown algal species.

**Figure S4.** Structural characteristics (% GC, %GC3, CDS size and intron number) of unbiased, gametophyte- and sporophyte-biased genes across brown algal species excluding orphan genes from the datasets.

**Figure S5. (A)** Evolutionary rates (dN/dS) for generation-biased genes when sex-biased genes were excluded from the analysis. GA: gametophyte, SP: sporophyte; SBG: sex-biased genes. **(B** and **C)** Comparison of evolutionary rates of unbiased genes with those of genes that exhibited either conserved bias (generation-biased in both species) or species-specific bias (i.e., were generation-biased in one species but unbiased in the other) in Ectocarpales **(B)** and Laminariales **(C)**. SBGs: sex-biased genes.

**Figure S6.** Non-synonymous substitutions **(A)**, synonymous substitutions **(B)** and codon usage bias **(C)** for unbiased, gametophyte- and sporophyte-biased genes in the four-studied species. Different letters above the plots indicate significant differences (P<0.05). Statistical significance was calculated by pairwise comparisons using the Wilcoxon rank sum test with Holm adjustment.

**Figure S7.** Evolutionary rates (dN/dS) in Laminariales and Ectocarpales orthologous genes with different evolutionary ages. Yellow dots indicate orthologous genes that are present in the four-studied species (‘old’ genes, i.e., ASOs), blue dots indicate order-specific orthologous genes (‘young’ genes, i.e. PSOs excluding ASOs). Overall, young genes evolve faster than older genes. Asterisks indicate a significant difference (Wilcoxon test, p-value<0.007).

